# Knockout of *ykcB*, a putative glycosyltransferase, leads to vancomycin resistance in *Bacillus subtilis*

**DOI:** 10.1101/2022.08.30.505962

**Authors:** Kazuya Ishikawa, Riko Shirakawa, Daiki Takano, Tomoki Kosaki, Kazuyuki Furuta, Chikara Kaito

## Abstract

Vancomycin resistance of gram-positive bacteria poses a serious health concern around the world. In this study, we searched for vancomycin-resistant mutants from a gene deletion library of a model gram-positive bacterium, *Bacillus subtilis*, to elucidate the mechanism of vancomycin resistance. We found that knockout of *ykcB*, a glycosyltransferase that is expected to utilize C55-P-glucose to glycosylate cell surface components, caused vancomycin resistance in *B. subtilis*. Knockout of *ykcB* altered the susceptibility to multiple antibiotics, including sensitization to β-lactams, and increased the pathogenicity to silkworms. Furthermore, the *ykcB*-knockout mutant had: i) an increased content of diglucosyl diacylglycerol, a glycolipid that shares a precursor with C55-P-glucose, ii) a decreased amount of lipoteichoic acid, and iii) decreased biofilm formation ability. These phenotypes and vancomycin resistance were abolished by knockout of *ykcC*, a *ykcB*-operon partner involved in C55-P-glucose synthesis. Overexpression of *ykcC* enhanced vancomycin resistance in both wild-type *B. subtilis* and the *ykcB*-knockout mutant. These findings suggest that *ykcB* deficiency induces structural changes of cell surface molecules depending on the *ykcC* function, leading to resistance to vancomycin, decreased biofilm formation ability, and increased pathogenicity to silkworms.

**IMPORTANCE:** Although vancomycin is effective against gram-positive bacteria, vancomycin-resistant bacteria is a major public health concern. While the vancomycin resistance mechanisms of clinically important bacteria such as *Staphylococcus aureus*, *Enterococcus faecium*, and *Streptococcus pneumoniae* are well-studied, they remain unclear in other gram-positive bacteria. In the present study, we searched for vancomycin-resistant mutants from a gene deletion library of a model gram-positive bacterium, *Bacillus subtilis*, and found that knockout of a putative glycosyltransferase, *ykcB*, caused vancomycin resistance in *B. subtilis*. Notably, unlike the previously reported vancomycin-resistant bacterial strains, *ykcB*-deficient *B. subtilis* exhibited increased virulence while maintaining its growth rate. Our results broaden the fundamental understanding of vancomycin-resistance mechanisms in gram-positive bacteria.

## INTRODUCTION

Antibiotics are widely used to treat bacterial infections, but the emergence of antibiotic-resistant bacteria is now a common and intractable problem. Vancomycin is a glycopeptide antibiotic that inhibits the polymerization of peptidoglycans in the cell wall of gram-positive bacteria by binding D-alanyl D-alanine residues. The emergence of vancomycin-resistant strains of *Staphylococcus aureus*, *Streptococcus pneumoniae*, and *Enterococcus faecium* poses a serious problem (1). In particular, vancomycin is one of the few effective antibiotics against methicillin-resistant *S. aureus* (MRSA), a bacterium resistant to many antibiotics, including β-lactams. Several cases of vancomycin-resistant MRSA have been reported (2). Understanding the mechanisms of vancomycin resistance is critical toward the development of new, effective antibiotics.

Vancomycin-resistant *S. aureus* is classified into 2 types: vancomycin-resistant *S. aureus* (VRSA) and vancomycin intermediate-resistant *S. aureus* (VISA) (2). While the vancomycin MIC of vancomycin-sensitive *S. aureus* is typically 0.5–2 μg/mL, the MICs for VRSA and VISA are ≥16 ug/ml and 4–8 ug/ml, respectively (3). The vancomycin resistance of VRSA is caused by acquisition of the *E. faecium vanA* gene, which encodes D-alanyl D-lactate ligase and changes the peptidoglycan structure (4). On the other hand, the vancomycin resistance of VISA is caused by gene mutations that result in cell wall thickening, such as *rpoB* (5), *graS* (6), *walK* (7), and *sdrC* (8). VRSA and VISA have reduced growth rates compared with vancomycin-sensitive *S. aureus* (8–11). The attenuation of virulence in VISA was demonstrated in a mouse model of sepsis and a *Galleria mellonella* infection model (12–14). VISA also has reduced ability to form biofilm, which is suggested to correlate with both the decreased virulence and vancomycin resistance (15, 16). While the molecular mechanisms underlying vancomycin resistance in clinically important bacteria, including *S. aureus*, have been studied, they remain unclear in other gram-positive bacteria.

In the present study, we searched for vancomycin-resistant mutants from a gene deletion library of a model gram-positive bacterium, *Bacillus subtilis*, and identified that knockout of *ykcB*, a putative glycosyltransferase, caused vancomycin resistance. *ykcB* is proposed to act cooperatively with *ykcC*, another glycosyltransferase, and *yngA*, a flippase on the plasma membrane (17). After *ykcC* produces the lipid phosphate carrier C55-P-glucose from UDP-glucose on the cytoplasmic side of the plasma membrane, C55-P-glucose is flipped to the outer surface of the plasma membrane by the function of *yngA*, and finally *ykcB* transfers a glucose from C55-P-glucose to some cell surface components (17). Here, we report that *ykcB* deficiency induces structural changes of the cell surface, such as a decreased amount of lipoteichoic acid, in a *ykcC*-dependent manner, leading to resistance to vancomycin.

## RESULTS

### Knockout of *ykcB* changes antibiotic susceptibility and increases virulence in silkworms

We searched for vancomycin-resistant strains among 3967 strains of the *B. subtilis* gene deletion library (18) and identified 23 strains with higher resistance to vancomycin than the parent strain (**Table 1**). Among the vancomycin-resistant strains, the *ykcB*-knockout mutant (Δ*ykcB*) exhibited the highest vancomycin resistance (**Fig. 1A**). To confirm that the vancomycin resistance was induced by *ykcB* knockout, we examined whether *ykcB* expression cancelled the vancomycin resistance of Δ*ykcB*. Introduction of *ykcB* into the *amyE* locus decreased vancomycin resistance in Δ*ykcB*, whereas introduction of empty vector into the *amyE* locus did not affect vancomycin resistance (**Fig. 1B**). These results suggest that loss of *ykcB* function leads to vancomycin resistance in *B. subtilis*.

**Table 1.**
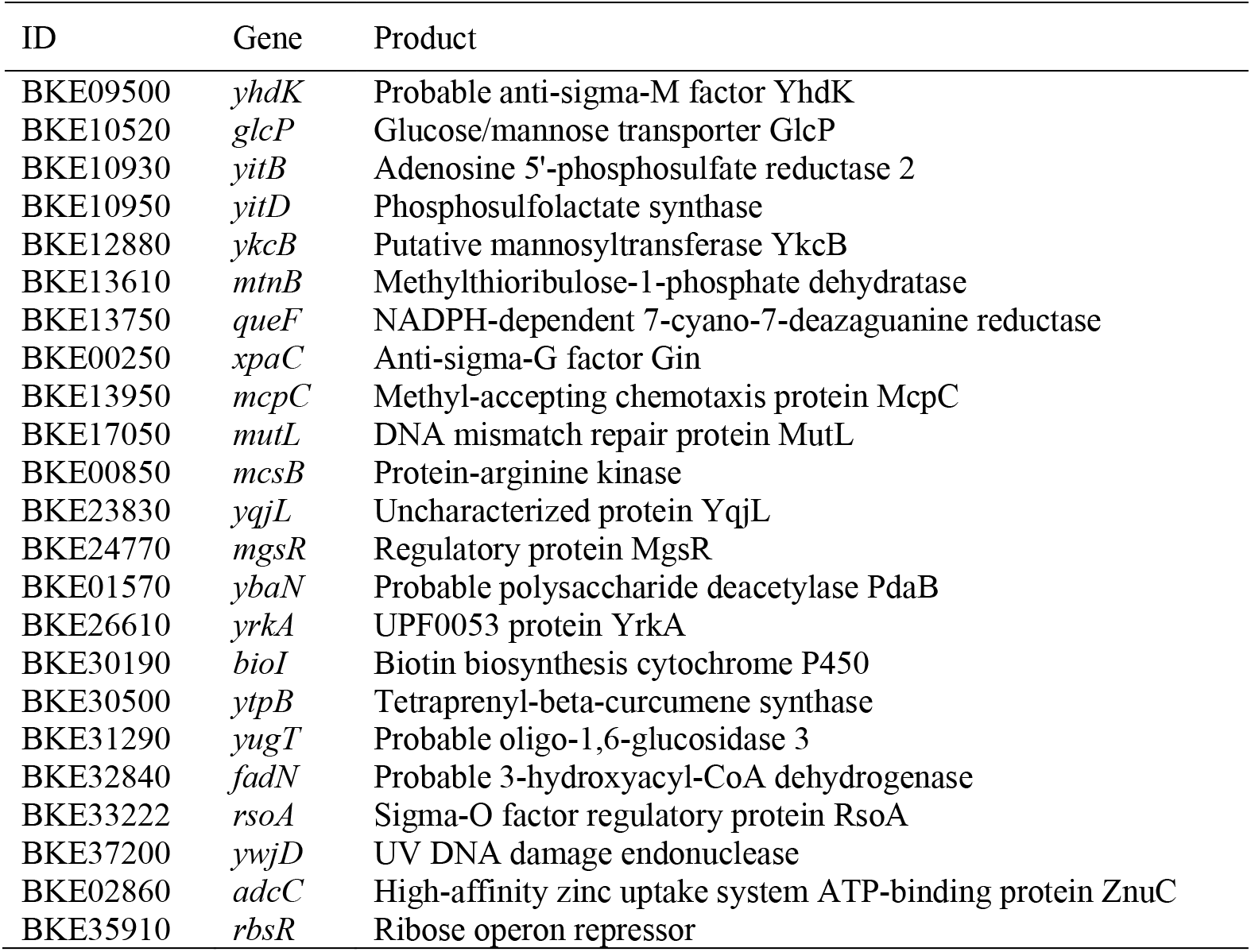
Gene knockout mutants resistant to vancomycin.

**Figure 1.**
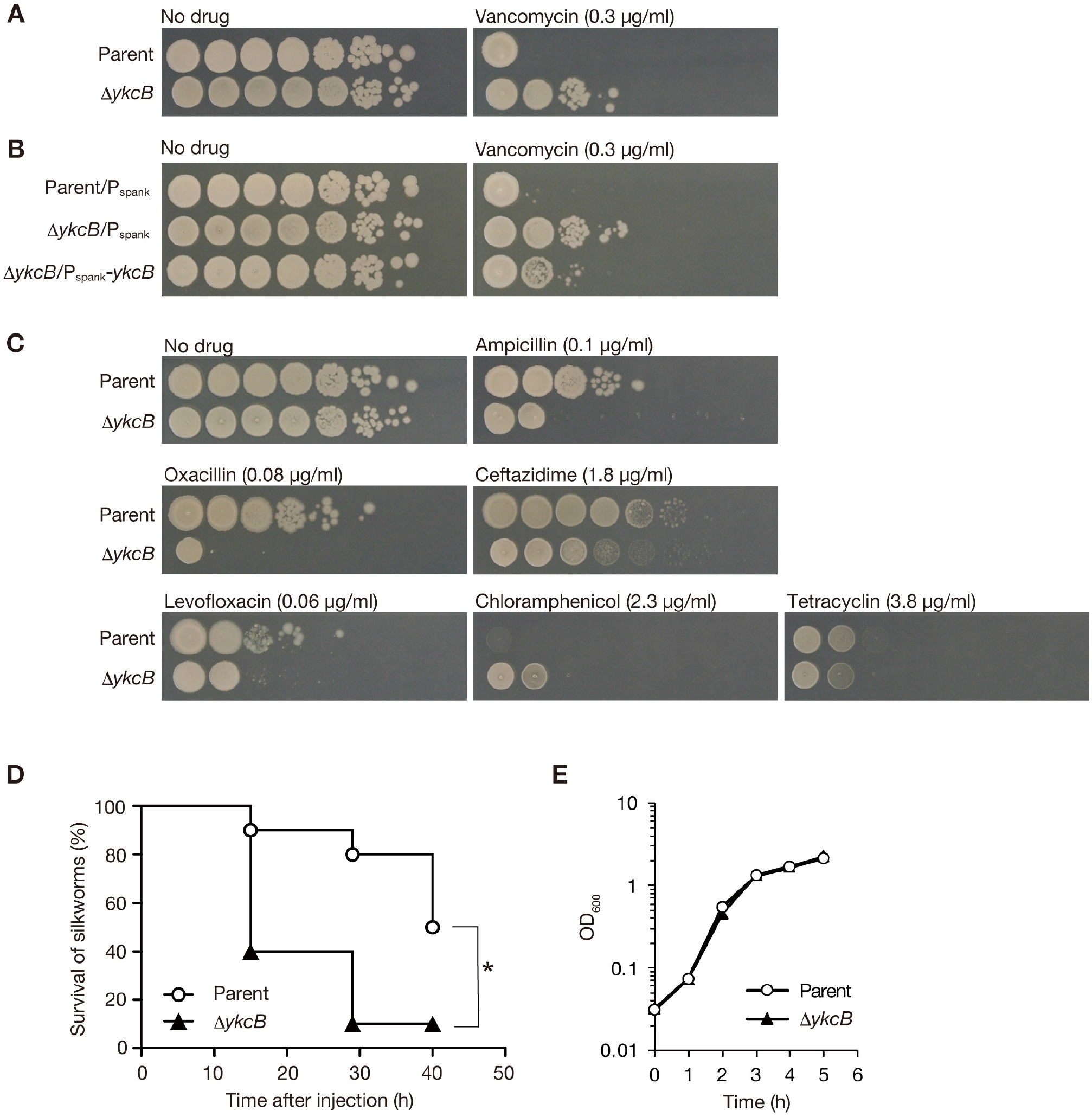
Knockout of *ykcB* alters sensitivity to antibiotics and increases silkworm killing activity. A. Overnight cultures of the parent strain (Parent) and *ykcB* knockout mutant (Δ*ykcB*) were serially diluted 10-fold and spotted onto LB plates supplemented with or without vancomycin (0.3 μg/ml). The plates were incubated overnight at 37°C. B. The parent strain transformed with empty vector (Parent/Pspank) and the *ykcB* knockout mutant transformed with empty vector (Δ*ykcB*/Pspank) or a vector encoding *ykcB* (Δ*ykcB*/Pspank-*ykcB*) were aerobically cultured overnight in the presence of 1 mM IPTG. The overnight cultures were serially diluted 10-fold and spotted onto LB plates supplemented with 1 mM IPTG and vancomycin or 1 mM IPTG alone. The plates were incubated overnight at 37°C. C. Overnight cultures of the parent strain (Parent) and the *ykcB* knockout mutant (Δ*ykcB*) were serially diluted 10-fold and spotted onto LB plates supplemented with or without ampicillin, oxacillin, ceftazidime, levofloxacin, chloramphenicol, or tetracycline. The plates were incubated overnight at 37°C. D. The silkworm killing activity of the parent strain (Parent) and *ykcB* knockout mutant (Δ*ykcB*) was examined. Silkworms (n=20) were injected with *B. subtilis* cells (8 × 10^6^ CFU) and silkworm survival was monitored. Asterisk indicates log-rank test p-value less than 0.05. E. The parent strain (Parent) and *ykcB* knockout mutant (Δ*ykcB*) were aerobically cultured in LB broth and the OD_600_ values of the cultures were measured.

We examined the sensitivity of Δ*ykcB* to antibiotics other than vancomycin. Δ*ykcB* became sensitive to the cell wall synthesis inhibitors ampicillin, oxacillin, and ceftazidime, and the DNA synthesis inhibitor levofloxacin (**Fig. 1C**). On the other hand, Δ*ykcB* became resistant to the protein synthesis inhibitor chloramphenicol and showed no change in sensitivity to tetracycline (**Fig. 1C**). These results indicate that *ykcB* deficiency alters the susceptibility to various antibiotics.

As vancomycin-resistant *S. aureus* strains are known to have attenuated pathogenicity (12–14), we investigated the pathogenicity of Δ*ykcB* using the silkworm infection model. Contrary to our expectation, silkworms injected with Δ*ykcB* died earlier than the parent strain, indicating increased virulence of Δ*ykcB* (**Fig. 1D**). Δ*ykcB* showed the same growth rate as the parent strain in nutrient medium (**Fig. 1E**), ruling out the possibility that the change in pathogenicity against silkworms depends on the bacterial growth rate.

### Vancomycin resistance in Δ*ykcB* is cancelled by the knockout of *ykcC*

To reveal more details of the mechanism of vancomycin resistance by *ykcB* knockout, we deleted the chromosomal region around the *ykcB* gene because *ykcB* and *ykcC* form an operon and are thought to be functionally related (19). Knockout of *mhqA* and *ykcC*, which respectively locate upstream and downstream of *ykcB*, did not lead to vancomycin resistance (**Fig. 2A, 2B**). Deletions of the intergenic region between *mhqA* and *ykcB* (*delA* and *delB*) caused slightly higher vancomycin resistance than that of the parent strain, but lower than that of Δ*ykcB* (**Fig. 2A, 2C**). Deletions in the *ykcB* coding region (*delC*, *delD*, *delE*, and *delF*) caused vancomycin resistance comparable to that of Δ*ykcB* (**Fig. 2A, 2C**). Introduction of a stop codon mutation in the *ykcB* gene in the *delA* background (*delA*/*ykcB*stop) caused a vancomycin-resistant phenotype to the same extent as Δ*ykcB*, but introduction of a stop codon mutation into both the *ykcB* and *ykcC* genes in the *delA* background (*delA*/*ykcB*stop/*ykcC*stop) did not cause a vancomycin-resistant phenotype, indicating that *ykcC* knockout cancelled the effect of *ykcB* knockout (**Fig. 2A, 2C**). These findings suggest that *ykcB* deficiency confers vancomycin resistance to *B. subtilis* in the presence of *ykcC*.

**Figure 2.**
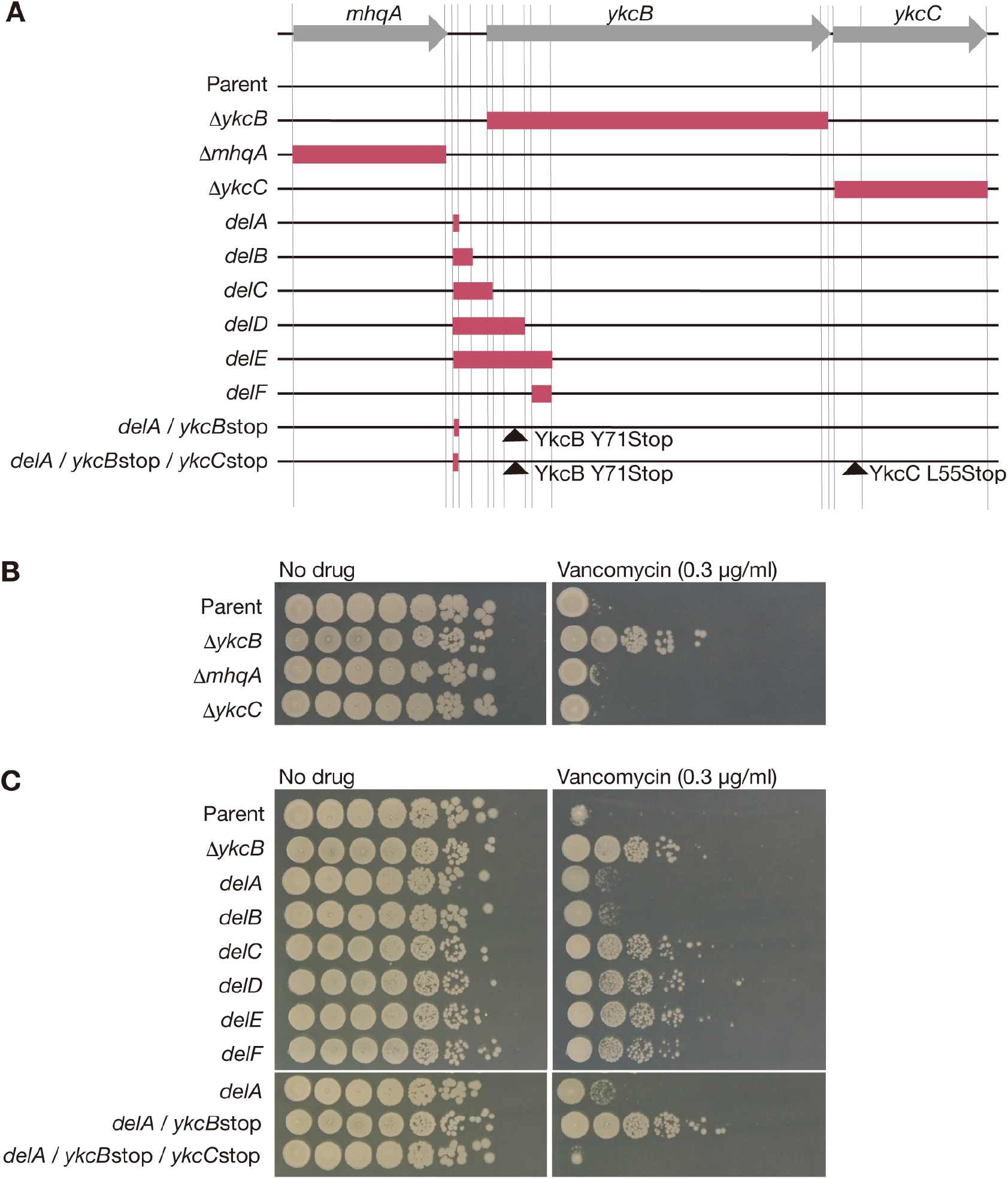
Knockout of *ykcB* leads to vancomycin resistance in a *ykcC-*dependent manner. A. Schematic representation of the *ykcB* flanking region is shown. The magenta box represents the chromosome region replaced with the erythromycin resistance gene in the gene knockout mutants or the chromosomal deletion mutants. Black arrowhead indicates the position at which the stop codon mutation was introduced. B. Overnight cultures of the parent strain (Parent), *ykcB* knockout mutant (Δ*ykcB*), *mhqA* knockout mutant (Δ*mhqA*), and *ykcC* knockout mutant (Δ*ykcC*) were serially diluted 10-fold and spotted onto LB plates supplemented with or without vancomycin. The plates were incubated overnight at 37°C. C. Overnight cultures of the parent strain (Parent), chromosomal deletion mutants, and stop codon mutants were serially diluted 10-fold and spotted onto LB plates supplemented with or without vancomycin. The plates were incubated overnight at 37°C.

### Knockout of *ykcB* increases the amount of diglucosyl diacylglycerol

According to the UniProt database, YkcB is predicted to be a glycosyltransferase with 14 transmembrane domains that belongs to the glycosyltransferase 39 family. In addition, it is assumed that YkcB utilizes C55-P-glucose, which is synthesized from UDP-glucose by YkcC, to glycosylate some cell surface molecules (17). Therefore, we hypothesized that *ykcB* knockout leads to the accumulation of C55-P-glucose and UDP-glucose, which results in an increased amount of diglucosyl diacylglycerol produced from UDP-glucose. Δ*ykcB* and the *delA*/*ykcB*stop mutants had increased amounts of diglucosyl diacylglycerol compared with the parent strain, indicating that the *ykcB* knockout increases the amount of diglucosyl diacylglycerol (**Fig. 3A, 3B**). In contrast, the *ykcC* knockout and the *delA*/*ykcB*stop/*ykcC*stop mutants did not have increased amounts of diglucosyl diacylglycerol (**Fig. 3A, 3B**), indicating that the *ykcB* knockout increases the amount of diglucosyl diacylglycerol in a *ykcC-*dependent manner.

**Figure 3.**
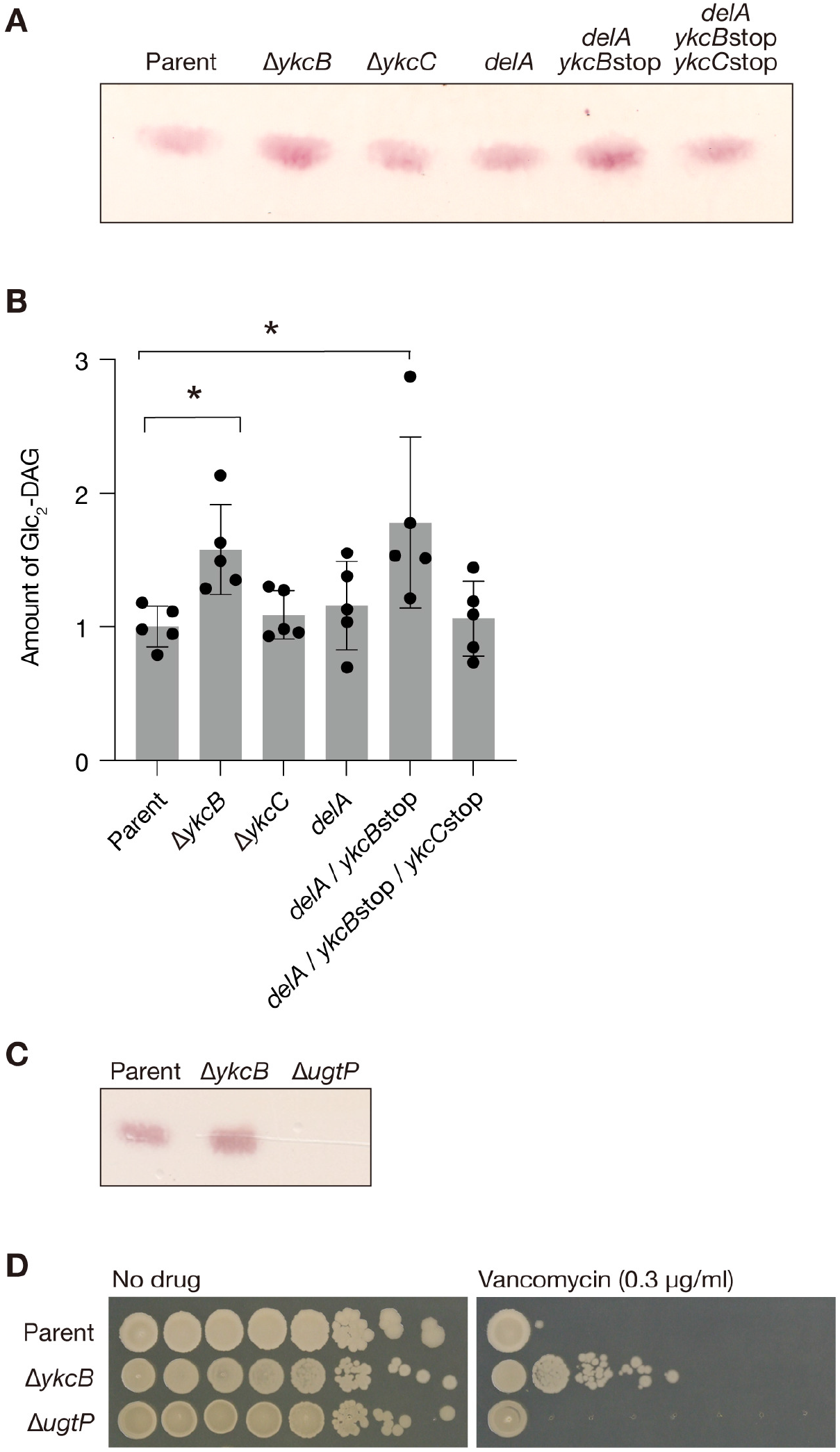
Knockout of *ykcB* leads to the accumulation of diglucosyl diacylglycerol in a *ykcC-*dependent manner. A. The *B. subtilis* parent strain (Parent), *ykcB* knockout mutant (Δ*ykcB*), *ykcC* knockout mutant (Δ*ykcC*), *delA* mutant (*delA*), *delA*/*ykcB*stop mutant (*delA*/*ykcB*stop), and *delA*/*ykcB*stop/*ykcC*stop mutant (*delA*/*ykcB*stop/*ykcC*stop) were cultured for 24 h and total lipids were extracted. Diglucosyl diacylglycerol was analyzed by TLC. B. The signal intensities of diglucosyl diacylglycerol in A were measured. Data are presented as means ± SD from 5 independent experiments. Stars indicate Dunnett’s multiple comparisons p value less than 0.05. C. The *B. subtilis* parent strain (Parent), *ykcB* knockout mutant (Δ*ykcB*), and *ugtP* knockout mutant (Δ*ugtP*) were cultured for 24 h and total lipids were extracted. Diglucosyl diacylglycerol was analyzed by TLC. D. Overnight bacterial cultures used in C were serially diluted 10-fold and spotted onto LB plates supplemented with or without vancomycin. The plates were incubated overnight at 37°C.

To determine whether diglucosyl diacylglycerol contributes to vancomycin resistance in *B. subtilis*, we examined the effect of knocking out the *ugtP* gene, which encodes a diglucosyl diacylglycerol synthetase. As expected, the *ugtP* knockout mutant did not produce diglucosyl diacylglycerol (**Fig. 3C**). The *ugtP* knockout mutant showed vancomycin sensitivity indistinguishable from that of the parent strain (**Fig. 3D**). Therefore, diglucosyl diacylglycerol does not contribute to vancomycin resistance.

### Knockout of *ykcB* decreases the amount of lipoteichoic acid and attenuates biofilm-forming ability

Based on the observation that Δ*ykcB* exhibits altered sensitivity to antibiotics, increased killing activity against silkworms, and an increased amount of diglucosyl diacylglycerol, we hypothesized that knockout of *ykcB* alters the amount of lipoteichoic acid or changes biofilm formation, both of which have important roles in antibiotic resistance and virulence. Δ*ykcB* and the *delA*/*ykcB*stop mutants had decreased amounts of lipoteichoic acid (**Fig. 4A, 4B**). Δ*ykcB* and the *delA*/*ykcB*stop mutants formed less biofilm than the parent strain and the *delA* mutant, respectively (**Fig. 5A, 5B**). These findings suggest that the *ykcB* knockout decreases the amount of lipoteichoic acid, and decreases biofilm forming ability. The *delA*/*ykcB*stop/*ykcC*stop mutant did not have a decreased amount of lipoteichoic acid or decreased biofilm formation (**Fig. 4A, 4B, 5A, 5B**), indicating that *ykcC* knockout cancelled the phenotypic changes caused by *ykcB* knockout.

**Figure 4.**
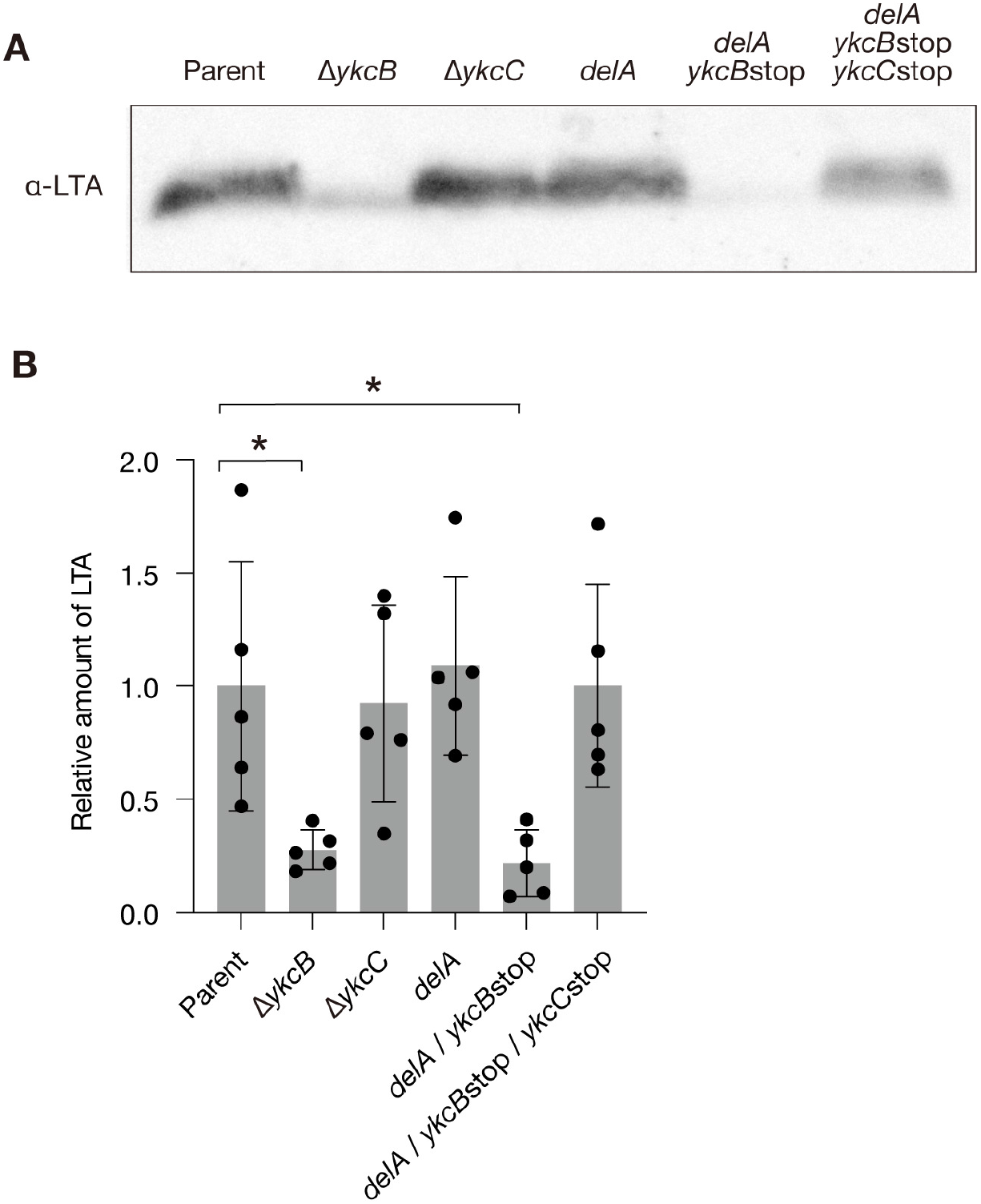
Knockout of *ykcB* decreases the amount of lipoteichoic acid in a *ykcC-*dependent manner. A. The *B. subtilis* parent strain (Parent), *ykcB* knockout mutant (Δ*ykcB*), *ykcC* knockout mutant (Δ*ykcC*), *delA* mutant (*delA*), *delA*/*ykcB*stop mutant (*delA*/*ykcB*stop), and *delA*/*ykcB*stop/*ykcC*stop mutant (*delA*/*ykcB*stop/*ykcC*stop) were cultured for 24 h and the lipoteichoic acids were extracted. Lipoteichoic acids were detected by Western blot analysis. B. The band intensities of lipoteichoic acids in A were measured. Data are presented as means ± SD from five independent experiments. Stars indicate Dunnett’s multiple comparisons p value less than 0.05.

**Figure 5.**
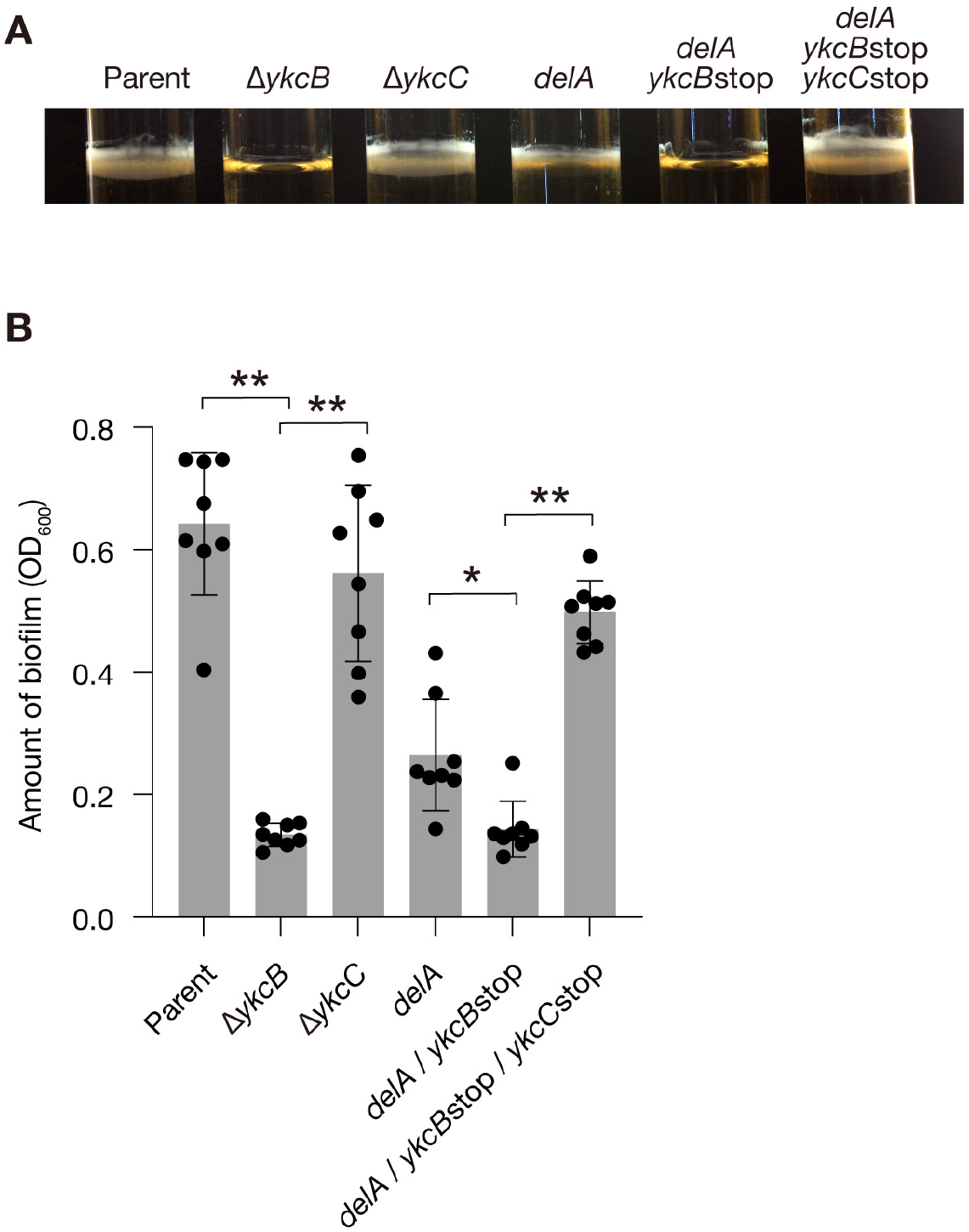
Knockout of *ykcB* decreases biofilm formation in a *ykcC*-dependent manner. A. The *B. subtilis* parent strain (Parent), *ykcB* knockout mutant (Δ*ykcB*), *ykcC* knockout mutant (Δ*ykcC*), *delA* mutant (*delA*), *delA*/*ykcB*stop mutant (*delA*/*ykcB*stop), and *delA*/*ykcB*stop/*ykcC*stop mutant (*delA*/*ykcB*stop/*ykcC*stop) were cultured for 2 days in glass tubes without shaking. The water surface areas were photographed. B. The amount of biofilm in A was measured. Data are presented as means ± SD from 8 independent experiments. Stars indicate Tukey’s multiple comparisons test p value less than 0.05.

### Overexpression of *ykcC* increases vancomycin resistance

Because *ykcC* knockout cancelled the vancomycin resistance caused by the *ykcB* knockout, expression of *ykcC* is hypothesized to have a positive role in vancomycin resistance. To evaluate this hypothesis, we transformed the parent and Δ*ykcB* strains with a multicopy plasmid encoding FLAG-tagged *ykcC* under the *ykcBC* native promoter. Western blot analysis revealed that the expression of FLAG-tagged *ykcC* was higher in Δ*ykcB* than in the parent strain (**Fig. 6A**), suggesting that positive feedback triggered by *ykcB* knockout upregulates the *ykcBC* promoter. The parent and Δ*ykcB* strains transformed with FLAG-tagged *ykcC* exhibited higher vancomycin resistance than those transformed with an empty vector (**Fig. 6B**). In addition, the Δ*ykcB* strain transformed with FLAG-tagged *ykcC* exhibited slightly higher vancomycin resistance than the parent strain transformed with FLAG-tagged *ykcC* (**Fig. 6B**). These findings suggest that *ykcC* confers vancomycin resistance to *B. subtilis* in an expression-dependent manner.

**Figure 6.**
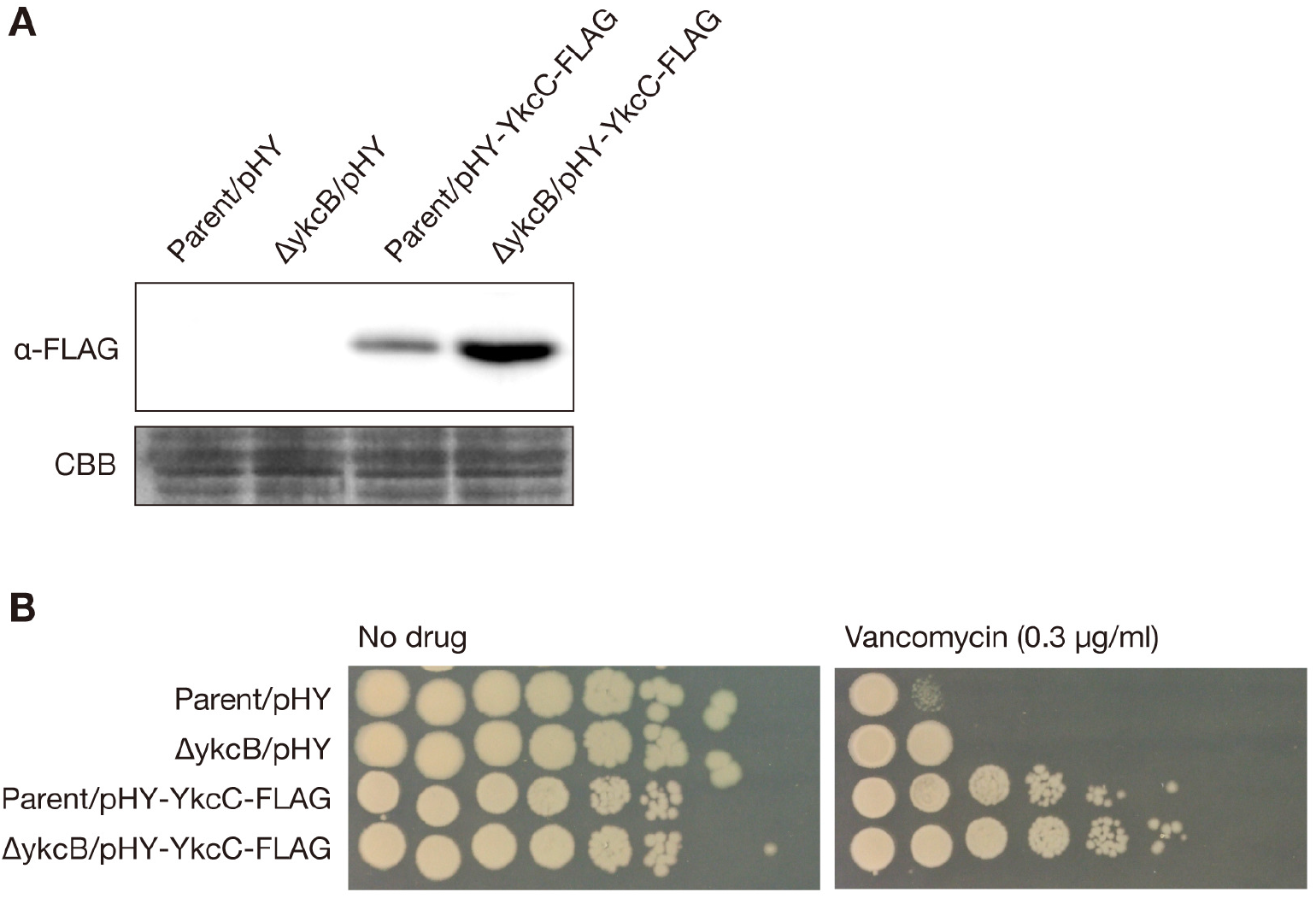
Overexpression of *ykcC* enhances vancomycin resistance in *B. subtilis*. A. The *B. subtilis* parent strain or the *ykcB* knockout mutant transformed with an empty vector (pHY) or a plasmid encoding FLAG-tagged *ykcC* (pHY-ykcC-FLAG) were subjected to Western blot analysis using the anti-FLAG antibody. The membrane was stained with Coomassie Brilliant Blue and shown as a loading control (CBB). B. Overnight bacterial cultures used in A were serially diluted 10-fold and spotted onto LB plates supplemented with or without vancomycin. The plates were incubated overnight at 37°C.

## DISCUSSION

The findings of the present study revealed that knockout of *ykcB*, a putative glycosyltransferase gene, confers *B. subtilis* resistance against vancomycin in a *ykcC*-dependent manner. Knockout of *ykcB* also leads to bacterial sensitivity to beta-lactams, decreases the amount of lipoteichoic acids, attenuates biofilm formation, and increases silkworm-killing activity. This study is the first to reveal that knockout of a specific gene leads to vancomycin resistance in *B. subtilis*.

The *ykcC* knockout mutant did not exhibit the same phenotypes as the *ykcB* knockout mutant. In addition, in the *ykcC*-stop codon mutant background, the stop codon mutation of *ykcB* led to no phenotypic changes. Therefore, the *ykcC* gene is required for the phenotypic changes triggered by *ykcB* knockout. Furthermore, overexpression of *ykcC* increases vancomycin resistance in the *B. subtilis* parent strain and the *ykcB* knockout strain. These findings suggest that expression of *ykcC* as well as knockout of *ykcB* increases the amount of some biologic molecule that leads to vancomycin resistance (**Fig. 7**). A previous study predicted that YkcC catalyzes UDP-glucose to C55-P-glucose and YkcB transfers glucose from C55-P-glucose to some cell surface molecule (17) (**Fig. 7**). Considering this prediction and the findings of the present study, C55-P-glucose might accumulate in the *ykcB*-knockout mutant and lead to various phenotypic changes, including vancomycin resistance (**Fig. 7**). C55-P-glucose is utilized as a sugar donor by YfhO to glycosylate lipoteichoic acid (17, 20). C55-P acts as an anchor to synthesize wall teichoic acid and peptidoglycan (21). Therefore, accumulation of C55-P-glucose might alter the structures of peptidoglycan, lipoteichoic acid, and wall teichoic acids, which may underlie the phenotypic changes observed in the *ykcB*-knockout mutant.

**Figure 7.**
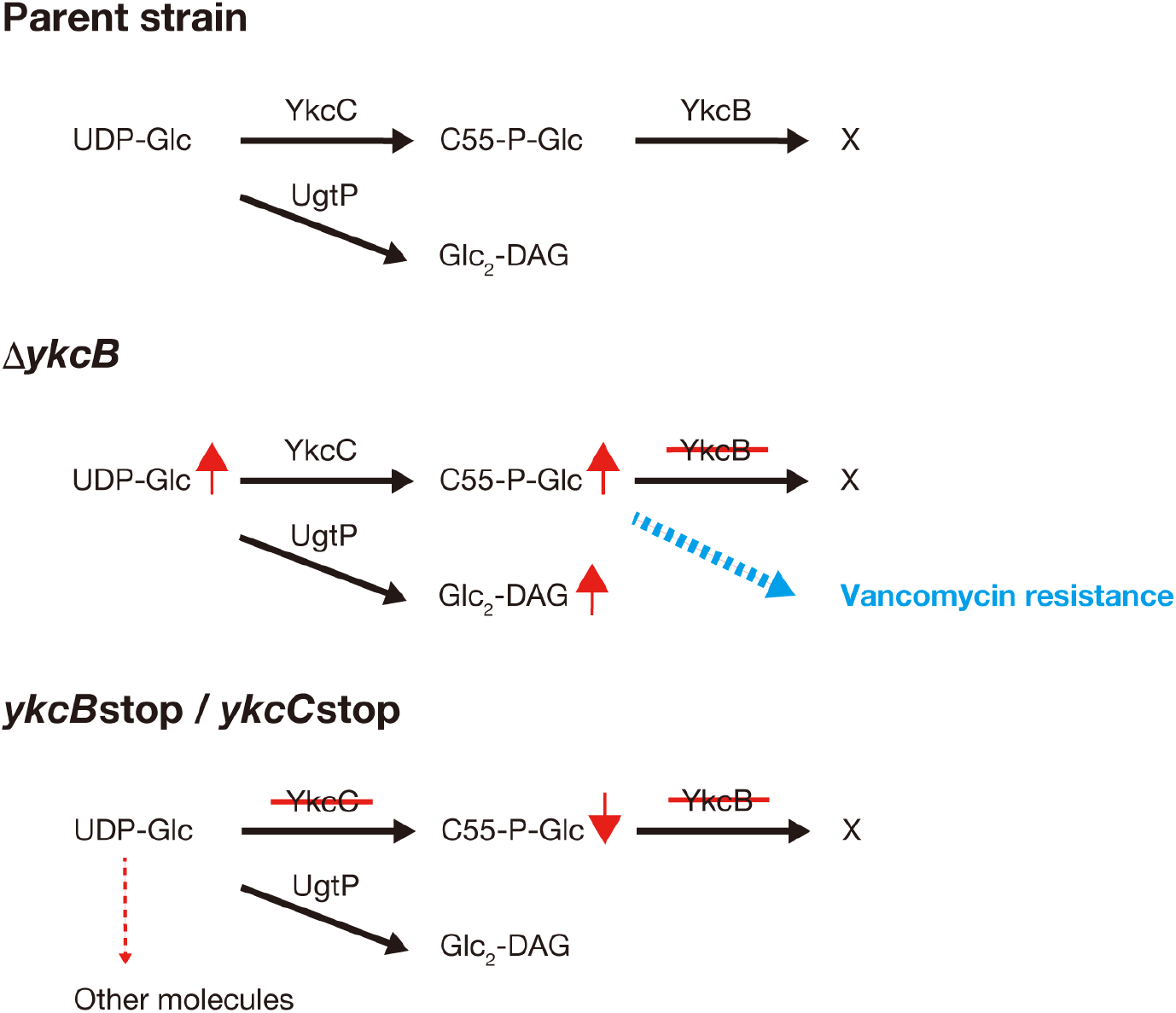
Model of vancomycin resistance induced by the *ykcB* knockout. Δ*ykcB* accumulates C55-P-glucose, which might change the cell surface structure and leads vancomycin resistance. In the *ykcB*stop/*ykcC*stop mutant, C55-P-glucose would not be accumulated because of the *ykcC* deficiency. The amount of diglucosyl diacylglycerol (Glc_2_-DAG) was not changed in the *ykcB*stop/*ykcC*stop mutant, suggesting that UDP-glucose is metabolized to other molecules in a *ykcC*-deficient background.

In the *ykcB*-knockout mutant, the amount of diglucosyl diacylglycerol was increased. We speculate that the accumulation of C55-P-glucose in the *ykcB-*knockout mutant increases UDP-glucose, a precursor of C55-P-glucose, and the increase in UDP-glucose leads to an increase in diglucosyl diacylglycerol (**Fig. 7**). Because knockout of *ugtP*, a synthetase gene of diglucosyl diacylglycerol, did not alter vancomycin resistance (**Fig. 3D**), the increased amount of diglucosyl diacylglycerol in the *ykcB*-knockout mutant does not contribute to the vancomycin resistance. In addition, in the *ykcC-*knockout mutant and the *ykcB*stop/*ykcC*stop mutant, diglucosyl diacylglycerol was not increased. In the absence of YkcC, UDP-glucose might be utilized for molecules other than diglucosyl diacylglycerol, which would prevent the accumulation of diglucosyl diacylglycerol (**Fig. 7**).

The *ykcB*-knockout mutant was resistant to vancomycin, but sensitive to beta-lactams. In VISA, a vancomycin resistance phenotype is accompanied by a beta-lactam sensitive phenotype, referred to as a “seesaw phenomenon” (22, 23). Thus, the beta-lactam sensitivity of the *ykcB*-knockout mutant of *B. subtilis* is consistent with that of VISA. The amounts of penicillin-binding protein 2 or phosphatidylglycerols are proposed to contribute to the beta-lactam sensitivity of VISA (24, 25). Further investigation is needed to examine penicillin-binding protein 2 and phosphatidylglycerols in the *ykcB-*knockout *B. subtilis* mutant.

This study demonstrated that the *ykcB-*knockout mutant has increased silkworm killing activity. In our previous study, *Escherichia coli* mutant strains resistant to vancomycin also showed resistance to antimicrobial peptides and increased silkworm-killing activity (26–28). The *B. subtilis ykcB-*knockout mutant might have resistance against silkworm antimicrobial peptides. In addition, a lipoteichoic acid synthetase gene knockout mutations in *Staphylococcus aureus* exhibits increased virulence against *Drosophila melanogaster* (29), leading to the proposal that lipoteichoic acid is a target molecule of Draper-dependent phagocytosis and lipoteichoic-deficient mutant bacteria escape the phagocytosis (29). The *ykcB-*knockout mutant might escape phagocytosis by silkworm immune cells because the *ykcB-*knockout mutant has little lipoteichoic acid.

In conclusion, this study identified that knockout of *ykcB* leads to vancomycin resistance in *B. subtilis*. The *ykcB-*knockout mutant exhibited increased virulence in silkworms, in contrast to VISA. Molecular investigation of vancomycin resistance using *B. subtilis*, a model gram-positive bacterium, is important to understand the conserved mechanism of vancomycin resistance between bacterial species.

## MATERIALS and METHODS

### Bacterial strains and culture conditions

*B. subtilis* 168 *trpC2* and its mutant strains were aerobically cultured in LB broth at 37°C. *B. subtilis* mutant strains carrying an erythromycin resistance gene were grown on LB plates containing erythromycin (1 μg/ml) and the colonies were aerobically cultured in LB broth without antibiotics at 37°C. *B. subtilis* strains transformed with pHY300PLK or pDR110 were cultured in LB broth containing tetracycline (30 μg/ml) or spectinomycin (50 μg/ml). Bacterial strains and plasmids used in this study are listed in **Table 2**.

**Table 2.**
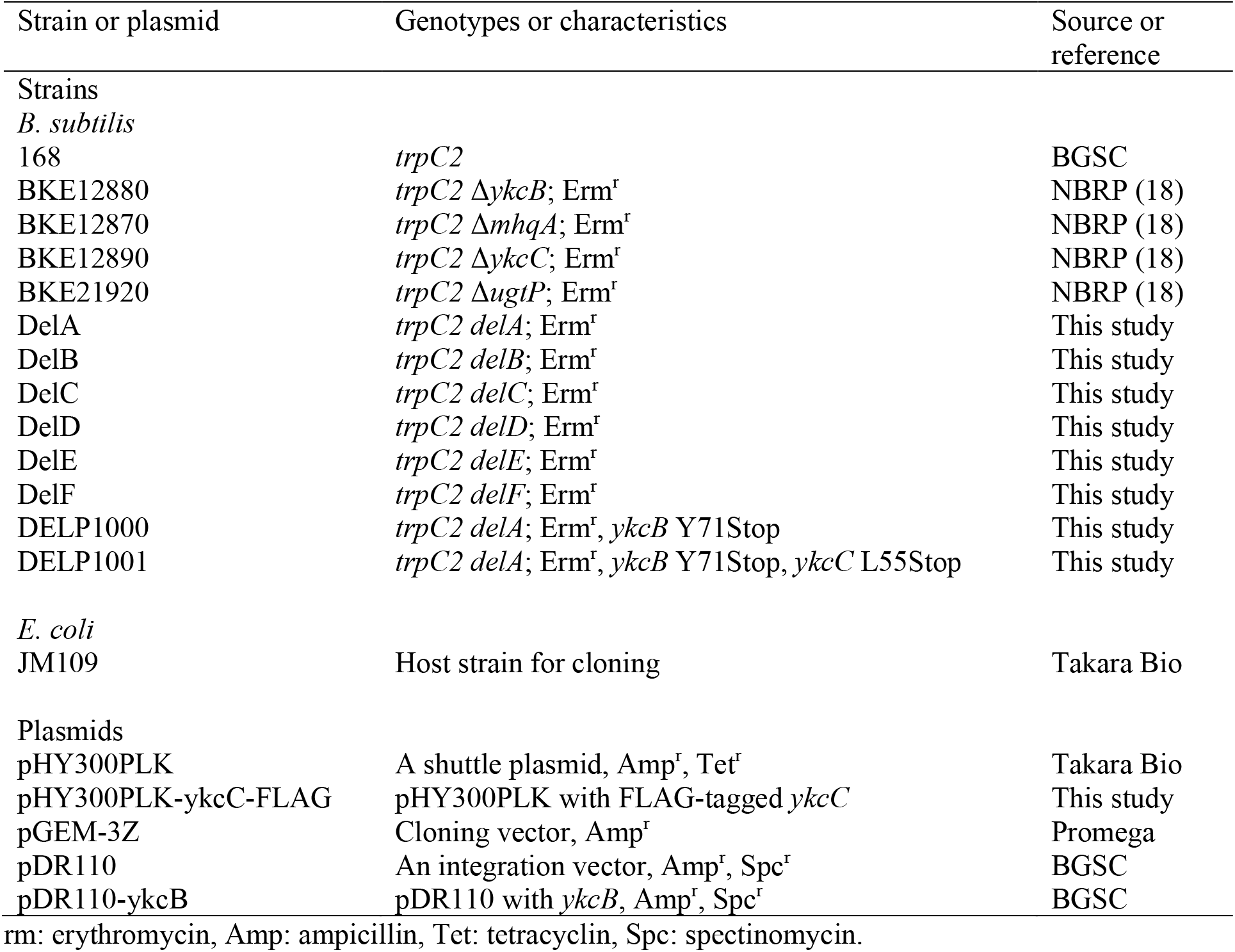
List of bacterial strains and plasmids used.

### Screening of vancomycin-resistant strains

The BKE library (18) was cultured in LB broth using a 96-well microplate at 37°C and the bacterial culture was spotted onto LB plates with or without vancomycin (0.45 μg/ml) using a replicator. The plates were incubated overnight at 37°C and mutant strains whose colonies appeared on vancomycin-containing plates were searched. The experiments were repeated and the strains were judged as vancomycin resistant when they formed colonies on vancomycin-containing plates in 2 experiments.

### Silkworm killing assay

Third instar silkworms were purchased from Ehime Sansyu (Ehime, Japan) and raised to fifth instar larvae by feeding them an artificial diet (Silkmate 2S; Nihon Nosan Kogyo Co., Kanagawa, Japan) at 27°C (30–32). The fifth instar hatched silkworms were fed an antibiotic-free artificial diet (Sysmex Co., Hyogo, Japan) for 1 day and used for infection experiments. *B. subtilis* overnight culture was diluted 5-fold with 0.9% saline and 0.05 ml was injected into the silkworm hemolymph using a tuberculin syringe equipped with 27-gauge needle. The OD_600_ values of *B. subtilis* overnight cultures were measured to confirm that the injected bacterial numbers were the same between strains.

### Genetic manipulation

#### 1) Construction of pDR110-ykcB

Genomic DNA of *B. subtilis* 168 *trpC2* was isolated using a QIAamp DNA blood minikit (Qiagen). A DNA fragment containing the *ykcB* gene was amplified by PCR using a 168 *trpC2* genomic DNA as a template and oligonucleotide primers (**Table 3**). The amplified DNA fragments were inserted into SphI and SalI sites in pDR110, resulting in pDR110-ykcB. Double crossover recombination of pDR110 or pDR110-ykcB at the *amyE* locus was confirmed by PCR using oligonucleotide primers (**Table 3**) and template genomic DNA from a spectinomycin-resistant colony.

**Table 3.**
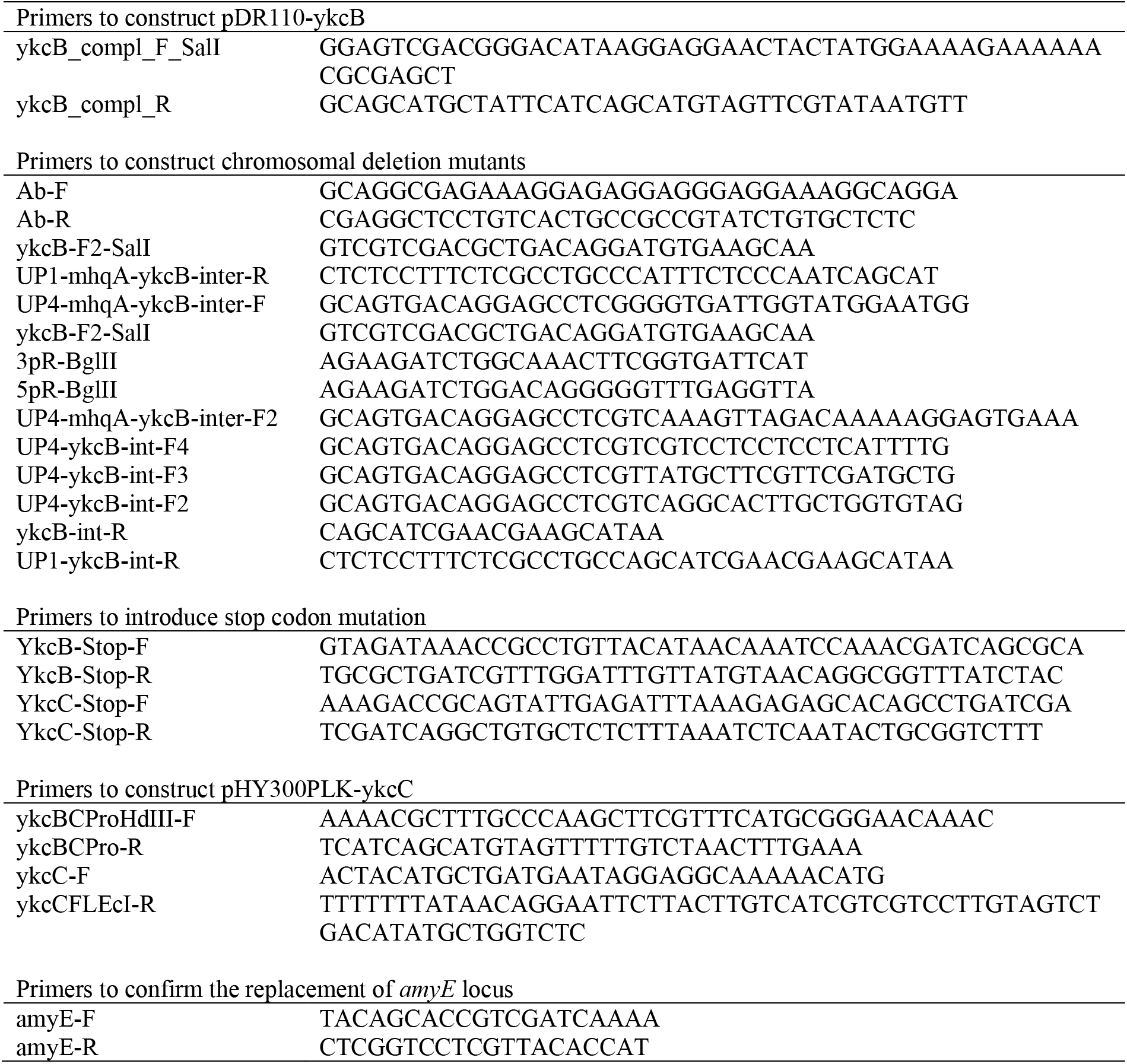
Primers used in this study.

#### 2) Construction of pHY300PLK-ykcC-FLAG

Two DNA fragments containing the promoter region of the *ykcBC* and *ykcC* ORF were amplified by PCR using oligonucleotide primers (**Table 3**) and the template genomic DNA of 168 *trpC2*. The 2 DNA fragments were connected by recombinant PCR and inserted into HindIII and EcoRI sites of pHY300PLK, resulting in pHY300PLK-ykcC-FLAG.

#### 3) Transformation by electroporation

Because the *ykcB* mutant did not have natural competency, we performed an electroporation to transform the *ykcB* mutant. *B. subtilis* overnight culture (1 ml) was inoculated into 100 ml of LB broth and aerobically cultured at 37°C until the OD_600_ reached 1.5. The culture was cooled on ice for 10 min and centrifuged at 3000 *g* for 10 min at 4°C. The bacterial pellet was suspended in ice-cold water. The washing procedure using ice-cold water was repeated 3 times and the bacterial pellet was suspended in 1 ml of 30% polyethylene glycol 6000. The bacterial suspension was frozen in liquid nitrogen and stored at −80°C. The frozen cells (100 μl) were thawed and mixed with plasmid DNA (200 ng). Electroporation (25 μF, 2500 V, 400 Ω) was performed in a 2-mm cuvette using the Gene Pulser Xcell Electroporation System (BioRad). After electroporation, the cells were immediately mixed with 2 ml SOC medium and incubated at 37°C for 90 min. The cells were spread onto LB plates containing appropriate selective antibiotics and incubated overnight at 37°C.

#### 4) Construction of chromosome deletion mutant by natural transformation

Targeting cassettes were constructed according to the previously described method (18) with minor modification. A DNA fragment containing the erythromycin resistance marker was amplified by PCR using oligonucleotide primers (**Table 3**) and a template genomic DNA from the *ykcB* mutant (BKE12880). The upstream and downstream DNA regions of the targeting chromosome locus were amplified by PCR using oligonucleotide primers (**Table 3**, **Table 4**) and a template genomic DNA from 168 *trpC2*. The 3 DNA fragments comprising the upstream and downstream regions and the erythromycin-resistance gene were mixed in an equal molar ratio and connected by PCR overlap extension using KOD FXneo DNA polymerase (Toyobo, Osaka, Japan). The connected DNA fragment was used for transformation without purification.

**Table 4.**
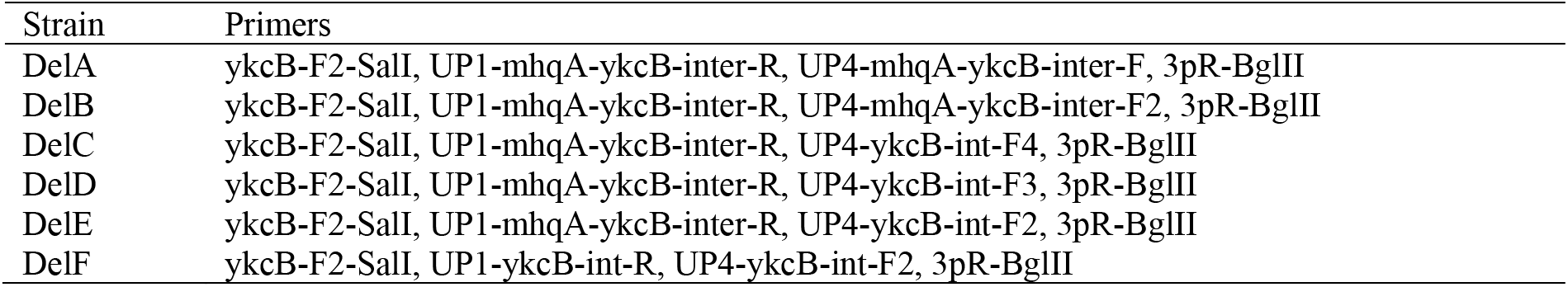
Primer sets used for constructing chromosomal deletion mutants.

| Strain | Primers |
| --- | --- |
| DelA | ykcB-F2-SalI, UP1-mhqA-ykcB-inter-R, UP4-mhqA-ykcB-inter-F, 3pR-BglII |
| DelB | ykcB-F2-SalI, UP1-mhqA-ykcB-inter-R, UP4-mhqA-ykcB-inter-F2, 3pR-BglII |
| DelC | ykcB-F2-SalI, UP1-mhqA-ykcB-inter-R, UP4-ykcB-int-F4, 3pR-BglII |
| DelD | ykcB-F2-SalI, UP1-mhqA-ykcB-inter-R, UP4-ykcB-int-F3, 3pR-BglII |
| DelE | ykcB-F2-SalI, UP1-mhqA-ykcB-inter-R, UP4-ykcB-int-F2, 3pR-BglII |
| DelF | ykcB-F2-SalI, UP1-ykcB-int-R, UP4-ykcB-int-F2, 3pR-BglII |

Competent cells for natural transformation were prepared according to the previous method (33) with minor modification. *B. subtilis* 168 *trpC2* overnight culture (50 μl) was inoculated into 5 ml of SPI medium (0.2% ammonium sulfate, 1.4% dipotassium hydrogen phosphate, 0.6% potassium dihydrogen phosphate, 0.1% trisodium citrate dihydrate, 0.02% magnesium sulfate heptahydrate, 0.5% glucose, 0.02% casamino acids, 0.1% yeast extract, 50 μg/ml L-leucine, 50 μg/ml L-methionine) and aerobically cultured at 37°C for 4.5 hour. Glycerol was added to the bacterial culture to a final concentration of 12.5%, frozen in a liquid nitrogen, and stored in a −80°C freezer. The frozen cells were thawed in a 37°C water bath and a 7.5-fold amount of SPII medium (0.2% ammonium sulfate, 1.4% dipotassium hydrogen phosphate, 0.6% potassium dihydrogen phosphate, 0.1% trisodium citrate dihydrate, 0.02% magnesium sulfate heptahydrate, 0.5% glucose, 5 mM magnesium chloride, 0.02% yeast extract, 5 μg/ml L-leucine, 5 μg/ml L-methionine) was added, and then the cells were aerobically cultured at 37°C for 90 min. A 50-μl amount of the cells was mixed with a targeting cassette and incubated at 37°C for 30 min. After adding 100 μl of LB broth to the cells, they were further incubated at 37°C for 60 min. The cells were spread onto LB plates containing 1 μg/ml erythromycin and incubated overnight at 37°C. The desired chromosomal deletion was confirmed by PCR.

#### 5) Construction of mutant strains carrying the stop codon mutation

A DNA fragment carrying the *mhqA-ykcBC* region and the erythromycin resistance gene was amplified by PCR using primer pairs (**Table 3**, ykcB-F2-SalI, 5pR-BglII) and template genomic DNA from the *delA* mutant. The DNA fragment was inserted into SalI and BglII sites of pGEM-3Z. Using the plasmid as a template, a thermal cycling reaction was performed using oligonucleotide primers to introduce the *ykcB* stop codon (**Table 3**). The reaction solution was digested with DpnI, and then used to transform the *E. coli* JM109 strain. A plasmid carrying the *ykcB* stop codon was purified from the *E. coli* colonies. Using the plasmid as a template, a thermal cycling reaction was performed using oligonucleotide primers to introduce the *ykcC* stop codon (**Table 3**), and the reaction solution was processed as described above. Plasmids carrying the *ykcB* stop codon and the *ykcC* stop codon were purified from the *E. coli* colonies. These 2 plasmids were digested with SalI and BglII and used for transformation of 168 *trpC2*. After transformation, genomic DNA was isolated from the erythromycin-resistant colonies and the desired stop codon mutations were confirmed by Sanger sequencing.

### Evaluation of antibiotic resistance

Autoclaved LB agar medium was mixed with antibiotic solutions and poured into a square dish (Eiken Chemical). *B. subtilis* overnight cultures were serially diluted 10-fold with LB broth in a 96-well microplate and 5 μl of the diluted bacterial solutions were spotted onto LB plates with or without antibiotics using an 8-channel Pipetman. The plates were incubated overnight at 37°C and photographed using a digital camera.

### Biofilm forming assay

*B. subtilis* overnight culture (20 μl) was inoculated into 2 ml of LB broth containing 1M NaCl in a glass tube and incubated for 2 days at 37°C. The bacterial culture containing biofilms was poured onto a Kimwipe placed on a Kimtowel (Nippon Paper Cresia, Tokyo, Japan). MilliQ water (2 ml) was added to the KimWipe, which was vortexed to detach the biofilms. The OD_600_ value of the solution was measured.

### Lipid extraction and TLC assay

*B. subtilis* overnight culture (1 ml) was added to 100 ml of LB broth and aerobically cultured at 37°C for 24 h; then, 40 ml of the bacterial culture was centrifuged at 10,400 *g* for 10 min at 4°C. The bacterial pellet was suspended with 1 ml of milliQ water and the lipids were extracted using the Bligh and Dyer method (34). The lipid fraction was evaporated by a centrifuge evaporator and the lipids were dissolved with 500 μl of chloroform:methanol (1:1 v/v). The sample was spotted onto TLC Silica gel 60 F_254_ (Merck) and the plate was developed in chloroform:methanol:water (65:25:4 v/v). Sugars were visualized by spraying a coloring agent (10.5 ml 15% 1-naphthol in ethanol, 40.5 ml ethanol, 6.5 ml sulfuric acid, and 4 ml water) and heating at 115°C.

### Western blot analysis

FLAG-tagged YkcC was detected according to a previous method (28) with minor modifications. *B. subtilis* overnight cultures was centrifuged at 21,400 *g* for 2 min and the bacterial pellet was frozen in liquid nitrogen. The bacterial pellet was thawed in buffer (50 mM Tris-HCl pH 7.8, 2 mM EDTA, 0.5 mM dithiothreitol, 0.4 mg/ml lysozyme) and subjected to freeze-thawing 2 times. TritonX-100 was added to the sample to produce a final concentration of 0.1% and the sample was incubated at 37°C for 30 min. An equal volume of 2x Laemmli sample buffer with 350 mM dithiothreitol was added to the sample and the sample was heated at 95°C for 3 min. The sample was centrifuged at 21500 *g* for 15 min, and the supernatant was electrophoresed in a 12% sodium dodecyl sulfate-polyacrylamide gel. Anti-DYKDDDDK (anti-FLAG) antibody (Wako, Japan) diluted 1:3000 in Canget signal solution 1 (Toyobo, Japan) was used as a first antibody solution. Anti-mouse IgG conjugated with horseradish peroxidase (HRP; Promega, Japan) diluted 1:3000 in Canget signal solution 2 (Toyobo, Japan) was used as a second antibody solution.

For detection of lipoteichoic acid, a previously described method (35) was used with modifications. *B. subtilis* overnight culture (50 μl) was inoculated to 5 ml of LB broth and aerobically cultured at 37°C for 24 h. The culture was centrifuged at 10,400 *g* for 10 min and the bacterial pellet was suspended in a 1.5x Laemmli sample buffer. The sample was boiled for 40 min and centrifuged at 10,400*g* for 10 min. The supernatants were electrophoresed in a 15% polyacrylamide gel and transferred to a nitrocellulose membrane (0.2 μm, Trans-Blot Transfer Medium, BioRad). The membrane was treated with 1:1000 anti- lipoteichoic acid antibody (clone 55, Hycult Biotech, Uden, The Netherlands) and washed 3 times with phosphate buffered saline. The membrane was treated with anti-mouse IgG HRP conjugate (Promega) and washed 3 times with phosphate buffered saline. The membrane was reacted with HRP substrate (Western Lightning, Perkin Elmer) and the signals were detected using ImageQuant LAS 4000 (Fujifilm, Tokyo, Japan). The band intensity was measured by Image J software (36).

### Statistical analysis

Survival curves of silkworms were analyzed by the log-rank test. The amounts of lipoteichoic acid and diglucosyl diacylglycerol were analyzed by Dunnett’s multiple comparisons test. The amount of biofilm was analyzed by Tukey’s multiple comparisons test. The statistical analysis was performed using Prism 9 (GraphPad Software).

## ACKNOWLEDGEMENT

This study was supported by JSPS Grants-in-Aid for Scientific Research (grants 22K14892, 22H02869, and 22K19435), the Takeda Science Foundation, the Ichiro Kanehara Foundation, and the Ryobi Teien Memory Foundation.

We thank the National BioResource Project-B. subtilis (National Institute of Genetics, Japan) for providing the *B. subtilis* BKE library and Bacillus Genetic Stock Center (BGSC) for providing the *B. subtilis* 168 *trpC2* and *B. subtilis* plasmids.

## REFERENCES

1. Karaman R, Jubeh B, Breijyeh Z. 2020. Resistance of Gram-Positive Bacteria to Current Antibacterial Agents and Overcoming Approaches. Molecules 25.

2. Gardete S, Tomasz A. 2014. Mechanisms of vancomycin resistance in Staphylococcus aureus. J Clin Invest 124:2836–40.

3. Swenson JM, Anderson KF, Lonsway DR, Thompson A, McAllister SK, Limbago BM, Carey RB, Tenover FC, Patel JB. 2009. Accuracy of commercial and reference susceptibility testing methods for detecting vancomycin-intermediate Staphylococcus aureus. J Clin Microbiol 47:2013–7.

4. Lessard IA, Walsh CT. 1999. VanX, a bacterial D-alanyl-D-alanine dipeptidase: resistance, immunity, or survival function? Proc Natl Acad Sci U S A 96:11028–32.

5. Matsuo M, Hishinuma T, Katayama Y, Cui L, Kapi M, Hiramatsu K. 2011. Mutation of RNA polymerase beta subunit (rpoB) promotes hVISA-to-VISA phenotypic conversion of strain Mu3. Antimicrob Agents Chemother 55:4188–95.

6. Cui L, Neoh HM, Shoji M, Hiramatsu K. 2009. Contribution of vraSR and graSR point mutations to vancomycin resistance in vancomycin-intermediate Staphylococcus aureus. Antimicrob Agents Chemother 53:1231–4.

7. Shoji M, Cui L, Iizuka R, Komoto A, Neoh HM, Watanabe Y, Hishinuma T, Hiramatsu K. 2011. walK and clpP mutations confer reduced vancomycin susceptibility in Staphylococcus aureus. Antimicrob Agents Chemother 55:3870–81.

8. Ishii K, Tabuchi F, Matsuo M, Tatsuno K, Sato T, Okazaki M, Hamamoto H, Matsumoto Y, Kaito C, Aoyagi T, Hiramatsu K, Kaku M, Moriya K, Sekimizu K. 2015. Phenotypic and genomic comparisons of highly vancomycin-resistant Staphylococcus aureus strains developed from multiple clinical MRSA strains by in vitro mutagenesis. Sci Rep 5:17092.

9. Foucault ML, Courvalin P, Grillot-Courvalin C. 2009. Fitness cost of VanA-type vancomycin resistance in methicillin-resistant Staphylococcus aureus. Antimicrob Agents Chemother 53:2354–9.

10. Pfeltz RF, Singh VK, Schmidt JL, Batten MA, Baranyk CS, Nadakavukaren MJ, Jayaswal RK, Wilkinson BJ. 2000. Characterization of passage-selected vancomycin-resistant Staphylococcus aureus strains of diverse parental backgrounds. Antimicrob Agents Chemother 44:294–303.

11. Sieradzki K, Tomasz A. 1997. Inhibition of cell wall turnover and autolysis by vancomycin in a highly vancomycin-resistant mutant of Staphylococcus aureus. J Bacteriol 179:2557–66.

12. Peleg AY, Monga D, Pillai S, Mylonakis E, Moellering RC, Jr., Eliopoulos GM. 2009. Reduced susceptibility to vancomycin influences pathogenicity in Staphylococcus aureus infection. J Infect Dis 199:532–6.

13. Howden BP, McEvoy CR, Allen DL, Chua K, Gao W, Harrison PF, Bell J, Coombs G, Bennett-Wood V, Porter JL, Robins-Browne R, Davies JK, Seemann T, Stinear TP. 2011. Evolution of multidrug resistance during Staphylococcus aureus infection involves mutation of the essential two component regulator WalKR. PLoS Pathog 7:e1002359.

14. Cameron DR, Ward DV, Kostoulias X, Howden BP, Moellering RC, Jr., Eliopoulos GM, Peleg AY. 2012. Serine/threonine phosphatase Stp1 contributes to reduced susceptibility to vancomycin and virulence in Staphylococcus aureus. J Infect Dis 205:1677–87.

15. Howden BP, Johnson PD, Ward PB, Stinear TP, Davies JK. 2006. Isolates with low-level vancomycin resistance associated with persistent methicillin-resistant Staphylococcus aureus bacteremia. Antimicrob Agents Chemother 50:3039–47.

16. Donlan RM, Costerton JW. 2002. Biofilms: survival mechanisms of clinically relevant microorganisms. Clin Microbiol Rev 15:167–93.

17. Wu CH, Rismondo J, Morgan RML, Shen Y, Loessner MJ, Larrouy-Maumus G, Freemont PS, Grundling A. 2021. Bacillus subtilis YngB contributes to wall teichoic acid glucosylation and glycolipid formation during anaerobic growth. J Biol Chem 296:100384.

18. Koo BM, Kritikos G, Farelli JD, Todor H, Tong K, Kimsey H, Wapinski I, Galardini M, Cabal A, Peters JM, Hachmann AB, Rudner DZ, Allen KN, Typas A, Gross CA. 2017. Construction and Analysis of Two Genome-Scale Deletion Libraries for Bacillus subtilis. Cell Syst 4:291–305 e7.

19. Ogura M, Ohsawa T, Tanaka T. 2008. Identification of the sequences recognized by the Bacillus subtilis response regulator YrkP. Biosci Biotechnol Biochem 72:186–96.

20. Rismondo J, Percy MG, Grundling A. 2018. Discovery of genes required for lipoteichoic acid glycosylation predicts two distinct mechanisms for wall teichoic acid glycosylation. J Biol Chem 293:3293–3306.

21. Kawai Y, Marles-Wright J, Cleverley RM, Emmins R, Ishikawa S, Kuwano M, Heinz N, Bui NK, Hoyland CN, Ogasawara N, Lewis RJ, Vollmer W, Daniel RA, Errington J. 2011. A widespread family of bacterial cell wall assembly proteins. EMBO J 30:4931–41.

22. Ortwine JK, Werth BJ, Sakoulas G, Rybak MJ. 2013. Reduced glycopeptide and lipopeptide susceptibility in Staphylococcus aureus and the “seesaw effect": Taking advantage of the back door left open? Drug Resist Updat 16:73–9.

23. Molina KC, Morrisette T, Miller MA, Huang V, Fish DN. 2020. The Emerging Role of beta-Lactams in the Treatment of Methicillin-Resistant Staphylococcus aureus Bloodstream Infections. Antimicrob Agents Chemother 64.

24. Werth BJ, Steed ME, Kaatz GW, Rybak MJ. 2013. Evaluation of ceftaroline activity against heteroresistant vancomycin-intermediate Staphylococcus aureus and vancomycin-intermediate methicillin-resistant S. aureus strains in an in vitro pharmacokinetic/pharmacodynamic model: exploring the “seesaw effect”. Antimicrob Agents Chemother 57:2664–8.

25. Hines KM, Shen T, Ashford NK, Waalkes A, Penewit K, Holmes EA, McLean K, Salipante SJ, Werth BJ, Xu L. 2020. Occurrence of cross-resistance and beta-lactam seesaw effect in glycopeptide-, lipopeptide- and lipoglycopeptide-resistant MRSA correlates with membrane phosphatidylglycerol levels. J Antimicrob Chemother 75:1182–1186.

26. Murakami K, Nasu H, Fujiwara T, Takatsu N, Yoshida N, Furuta K, Kaito C. 2021. The Absence of Osmoregulated Periplasmic Glucan Confers Antimicrobial Resistance and Increases Virulence in Escherichia coli. J Bacteriol 203:e0051520.

27. Kaito C, Yoshikai H, Wakamatsu A, Miyashita A, Matsumoto Y, Fujiyuki T, Kato M, Ogura Y, Hayashi T, Isogai T, Sekimizu K. 2020. Non-pathogenic Escherichia coli acquires virulence by mutating a growth-essential LPS transporter. PLoS Pathog 16:e1008469.

28. Nasu H, Shirakawa R, Furuta K, Kaito C. 2022. Knockout of mlaA increases Escherichia coli virulence in a silkworm infection model. PLoS One 17:e0270166.

29. Hashimoto Y, Tabuchi Y, Sakurai K, Kutsuna M, Kurokawa K, Awasaki T, Sekimizu K, Nakanishi Y, Shiratsuchi A. 2009. Identification of lipoteichoic acid as a ligand for draper in the phagocytosis of Staphylococcus aureus by Drosophila hemocytes. J Immunol 183:7451–60.

30. Kaito C, Murakami K, Imai L, Furuta K. 2020. Animal infection models using non-mammals. Microbiol Immunol 64:585–592.

31. Kaito C, Akimitsu N, Watanabe H, Sekimizu K. 2002. Silkworm larvae as an animal model of bacterial infection pathogenic to humans. Microb Pathog 32:183–90.

32. Kaito C, Kurokawa K, Matsumoto Y, Terao Y, Kawabata S, Hamada S, Sekimizu K. 2005. Silkworm pathogenic bacteria infection model for identification of novel virulence genes. Mol Microbiol 56:934–44.

33. Sadaie Y, Kada T. 1983. Formation of competent Bacillus subtilis cells. J Bacteriol 153:813–21.

34. Bligh EG, Dyer WJ. 1959. A rapid method of total lipid extraction and purification. Can J Biochem Physiol 37:911–7.

35. Garcia-Gomez E, Miranda-Ozuna JFT, Diaz-Cedillo F, Vazquez-Sanchez EA, Rodriguez-Martinez S, Jan-Roblero J, Cancino-Diaz ME, Cancino-Diaz JC. 2017. Staphylococcus epidermidis lipoteichoic acid: exocellular release and ltaS gene expression in clinical and commensal isolates. J Med Microbiol 66:864–873.

36. Schneider CA, Rasband WS, Eliceiri KW. 2012. NIH Image to ImageJ: 25 years of image analysis. Nat Methods 9:671–5.

